# Predicting metal-binding proteins and structures through integration of evolutionary-scale and physics-based modeling

**DOI:** 10.1101/2024.08.09.607368

**Authors:** Xin Dai, Max Henderson, Shinjae Yoo, Qun Liu

## Abstract

Metals are essential elements in all living organisms, binding to approximately 50% of proteins. They serve to stabilize proteins, catalyze reactions, regulate activities, and fulfill various physiological and pathological functions. While there have been many advancements in determining the structures of protein-metal complexes, numerous metal-binding proteins still need to be identified through computational methods and validated through experiments. To address this need, we have developed the ESMBind workflow, which combines evolutionary scale modeling (ESM) for metal-binding prediction and physics-based protein-metal modeling. Our approach utilizes the ESM-2 and ESM-IF models to predict metal-binding probability at the residue level. In addition, we have designed a metal-placement method and energy minimization technique to generate detailed 3D structures of protein-metal complexes. Our workflow outperforms other models in terms of residue and 3D-level predictions. To demonstrate its effectiveness, we applied the workflow to 142 uncharacterized fungal pathogen proteins and predicted metal-binding proteins involved in fungal infection and virulence.

## Introduction

Protein-metal ion interactions are fundamental to a wide range of biological processes, including enzyme catalysis, signal transduction, structural stabilization, and the regulation of various physiological and pathological activities. These interactions exhibit distinct patterns in coordination geometry and binding specificity that reflect their chemical nature. It is estimated that around 50% of proteins bind metals ^1^. Gaining insights into these interactions at the atomic level is critical for understanding protein function and holds promise for applications in drug discovery and protein engineering. However, accurately predicting metal-binding sites and the precise locations of metals within 3D protein structures remains a significant challenge, particularly given the experimental uncertainties in structure determination and the complexity of metal coordination geometries^2^.

Recent advances in deep learning have unlocked new possibilities for improving metal-binding site prediction by utilizing large-scale protein data^3–7^. In particular, pretrained foundation models like ESM-2 and ESM-IF have emerged as powerful tools for understanding protein sequences and structures. ESM-2 is a large language model trained on approximately 65 million unique sequences from the UniRef database using a masked language modeling objective. By learning to predict masked amino acids from their surrounding context, ESM-2 captures sequence patterns across evolutionarily diverse proteins and implicitly learns structural information, despite being trained solely on sequence data. ESM-IF complements this sequence-based approach by directly modeling the relationship between protein sequences and structures. Trained on 12 million protein structures predicted by AlphaFold-2^8^, ESM-IF combines a transformer architecture with invariant geometric processing layers to predict protein sequences from backbone atom coordinates. Its high native sequence recovery rates, particularly for buried residues, demonstrate its deep understanding of sequence-structure relationships. These pretrained models can be fine-tuned or used as feature extractors for downstream tasks, often outperforming models trained from scratch on smaller, task-specific datasets.

Deep learning approaches for metal binding prediction operate at three distinct levels: sequence-level, residue-level, and atomic-level modeling. Sequence-level prediction aims to determine whether a protein sequence will bind metal ions at all, providing a binary or probabilistic classification for metal-binding proteins^9^. While this level of prediction is valuable for proteome-wide annotation and functional studies, in this work we focus on the more detailed residue-level and atomic-level predictions.

At the residue level, methods like LMetalSite^10^ and M-Ionic^11^ identify which specific amino acid residues are likely to participate in metal binding. LMetalSite utilizes ProtTrans^3^, a pretrained language model similar to ESM-2, while M-Ionic directly employs ESM-2 to predict metal-binding residues. While these approaches effectively identify potential binding residues for various metal ions, they do not address the subsequent challenge of determining the precise 3D coordinates of the metal ions within the protein structure.

The transition from residue-level prediction to atomic-level modeling presents several significant challenges in protein-metal binding prediction. First, deep learning models often predict high probabilities for multiple metal types at the same binding sites, making it difficult to determine the specific metal present. Second, even with accurate binding residue predictions, determining the number of metal ions and their precise 3D coordinates remains complex. These challenges require a solution that integrates both geometric constraints and energetic considerations while maintaining physically realistic coordination geometry.

Current atomic-level modeling approaches address these challenges with varying success. Metal3D^12^ approaches this challenge using 3D convolutional neural networks, achieving high spatial accuracy for zinc placement with predictions within 0.70 ± 0.64 Å of experimental positions. However, Metal3D’s capability is currently limited to zinc, leaving the prediction of other metal types and their precise positioning as an open challenge. Taking a different approach to atomic-level modeling, AlphaFill^13^ enriches AlphaFold-predicted structures by transplanting metal ions and other small molecules from experimentally determined structures based on sequence and structural homology. While this method can accurately place metals in proteins with known homologs, it struggles with novel metal-binding sites or proteins lacking close structural relatives in experimental databases.

In this study, we address these challenges by developing ESMBind, a novel workflow that bridges the gap between residue-level prediction and atomic-level modeling for seven essential-to-life metals: Zn^2+^, Ca^2+^, Mg^2+^, Mn^2+^, Fe^3+^, Co^2+^, and Cu^2+^. ESMBind combines the power of deep learning with physics-based modeling through a two-stage approach. In the first stage, it integrates sequence information from the pretrained ESM-2 model and structural information from ESM-IF to predict metal binding probabilities at the residue level. The second stage transforms these residue-level predictions into precise 3D coordinates through a sophisticated local grid search strategy and energy minimization procedure. The key innovation of ESMBind lies in its comprehensive approach to metal placement. Our local grid search strategy systematically explores possible metal positions while accounting for the number of ions present in the structure. This initial placement is followed by energy minimization that optimizes the positions of metal ions while preserving physically realistic coordination geometry. This combined approach ensures both geometric accuracy and energetic feasibility in the final protein-metal complex structures. We trained and evaluated our approach using high-quality protein-metal binding data from the BioLip database^14^, which curates information from the Protein Data Bank (PDB)^15^. By comparing our predictions with the ground truth data, we demonstrate the accuracy and effectiveness of our method in both identifying protein-metal binding sites and determining the optimal positions of metal ions within protein structures.

## Results

### Overview of the workflow

Our ESMBind workflow is illustrated in Figure 1a. This workflow integrates deep learning-based prediction with physics-based modeling to enhance prediction accuracy of protein-metal binding sites. The workflow is designed to predict binding sites for seven common metal ions in metalloproteins: Zn^2+^, Ca^2+^, Mg^2+^, Mn^2+^, Fe^3+^, Co^2+^, and Cu^2+^.

**Figure 1.**
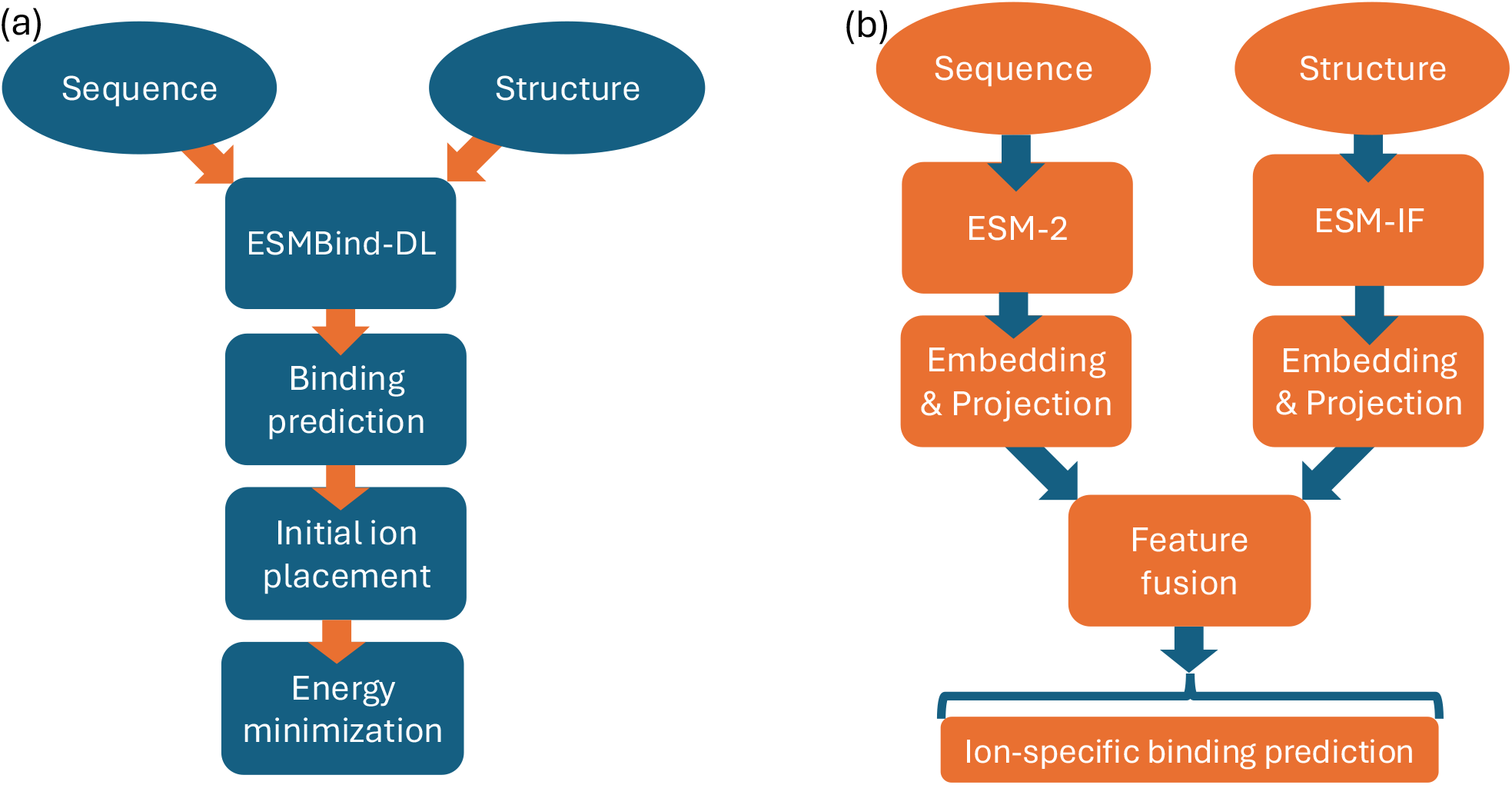
ESMBind workflow and architecture. (a) Overview of the ESMBind workflow for metal-binding site prediction. The process begins with protein sequence and structure as inputs. These are fed into the ESMBind-DL model (detailed in panel b), which predicts binding probabilities at the residue level. These probabilities guide initial ion placement, followed by energy minimization to produce the final 3D structure with placed ions. (b) Architecture of the ESMBind-DL model. The model processes protein sequence and structure information through ESM-2 and ESM-IF, respectively. The resulting sequence and structure embeddings undergo projection to reduce dimensionality. These projected features are then combined through feature fusion. The fused features are processed by individual ion prediction heads, each generating binding probabilities for a specific metal type at the residue level.

The workflow begins with the input of protein sequence and structure data. ESMBind processes the input information to predict binding probabilities for each residue to each of the seven metals, performing a binary classification at the residue level to determine whether a residue binds to a metal ion. Following this residue-level prediction, the workflow proceeds to the metal placement stage. Here, the binding probabilities are used to guide the initial placement of metals within the protein structure. This step transforms the residue-level predictions into a 3D structure of the protein-metal complex. The final step involves energy minimization, which optimizes the protein structure and metal positions according to physical principles using the AMBER force field. This step optimizes the initial ion placements, ensuring they are physically and stereochemically plausible within the context of the protein structure. The output of this stage is a 3D structure of the protein modeled with a predicted metal-binding site.

### Model architecture

The ESMBind-DL model is a deep learning architecture designed to predict protein-metal binding probabilities at the residue level. It leverages embeddings from two pretrained models: ESM-2 and ESM-IF. ESM-2 captures sequence information, while ESM-IF, trained on 3D structures, provides structural context.

As illustrated in Figure 1b, the model processes sequence input through ESM-2 (650M variant), generating 1280-dimensional embeddings per residue, and structural input through ESM-IF, yielding 512-dimensional embeddings per residue. These embeddings are then processed by separate feature processing modules. Each module consists of layer normalization, dropout, a linear layer, and a LeakyReLU activation function. The feature processing modules reduce the ESM-2 and ESM-IF embeddings to 256 dimensions each. These processed embeddings are then concatenated to form a 512-dimensional unified representation. This combined vector undergoes another feature processing module, transforming it into a 128-dimensional feature vector that integrates both sequence and structure information. Finally, the model feeds this feature vector into metal-specific classifier heads. Each head is responsible for predicting the binding probability of a particular metal ion at the residue level. This architecture allows ESMBind-DL to simultaneously predict binding events for multiple metal types, leveraging both sequence and structural data.

### Residue-level prediction

The residue-level prediction forms the foundation of ESMBind-DL’s capabilities. For each metal type, the model predicts the probability of each residue in the protein sequence binding to that metal. This task is formulated as a binary classification problem, where each residue is classified as either binding or non-binding for a specific metal type.

As shown in Table 2, the dataset exhibits substantial class imbalance, with binding residues accounting for only 0.91-1.53% of total residues across different metal types. Given this imbalance, we selected the Area Under the Precision-Recall Curve (AUPRC) as our primary evaluation metric, supplemented by precision, recall, F1 score, and Area Under the ROC Curve (AUC) for a comprehensive assessment. Table 1 presents a comparison between ESMBind-DL and two existing methods, M-Ionic and LMetalSite, with results ordered by test set size. For Zn^2+^, which has the largest test set, ESMBind-DL outperforms both existing methods in AUPRC. ESMBind-DL and LMetalSite show comparable performance on the test datasets of similar sizes for Mg^2+^ and Ca^2+^. For metals with medium-sized test sets, LMetalSite achieves the highest AUPRC on Mn^2+^, while ESMBind-DL demonstrates superior performance on Fe^3+^ compared to M-Ionic. ESMBind-DL maintains its effectiveness even for metals with the smallest test sets (Co^2+^ and Cu^2+^), consistently outperforming M-Ionic. Notably, LMetalSite was not trained on Fe^3+^, Co^2+^, and Cu^2+^, thus comparisons for these metals are unavailable. These results demonstrate that ESMBind-DL achieves robust performance across different metal types, regardless of test set size. This reliable residue-level prediction provides a solid foundation for the subsequent steps in the workflow, namely the initial metal placement and energy minimization.

**Table 1.** Performance comparison of ESMBind-DL with state-of-the-art methods on metal ion binding site prediction (bold: highest values among compared methods).

| Data set | Method | Rec | Pre | F1 | AUC | AUPRC |
| --- | --- | --- | --- | --- | --- | --- |
| $\text{Zn}^{2+}$ (420) | ESMBind-DL | 0.637 | **0.832** | **0.722** | 0.982 | **0.763** |
|  | M-Ionic | **0.730** | 0.667 | 0.697 | 0.981 | 0.747 |
|  | LMetalSite | 0.515 | 0.634 | 0.568 | **0.985** | 0.608 |
| $\text{Mg}^{2+}$ (408) | ESMBind-DL | 0.303 | **0.597** | 0.402 | 0.875 | 0.343 |
|  | M-Ionic | 0.339 | 0.404 | 0.369 | 0.845 | 0.307 |
|  | LMetalSite | **0.376** | 0.452 | **0.411** | **0.906** | **0.344** |
| $\text{Ca}^{2+}$ (361) | ESMBind-DL | 0.326 | **0.752** | **0.455** | 0.892 | **0.449** |
|  | M-Ionic | **0.442** | 0.427 | 0.434 | 0.884 | 0.430 |
|  | LMetalSite | 0.382 | 0.492 | 0.430 | **0.902** | 0.398 |
| $\text{Mn}^{2+}$ (121) | ESMBind-DL | 0.585 | **0.818** | 0.682 | 0.981 | 0.754 |
|  | M-Ionic | 0.591 | 0.741 | 0.658 | 0.975 | 0.676 |
|  | LMetalSite | **0.750** | 0.651 | **0.697** | **0.990** | **0.773** |
| $\text{Fe}^{3+}$ (58) | ESMBind-DL | **0.723** | 0.810 | **0.764** | 0.974 | **0.800** |
|  | M-Ionic | 0.657 | **0.815** | 0.728 | 0.974 | 0.745 |
| $\text{Co}^{2+}$ (51) | ESMBind-DL | **0.526** | 0.741 | **0.615** | 0.926 | **0.569** |
|  | M-Ionic | 0.179 | **0.791** | 0.292 | **0.937** | 0.531 |
| $\text{Cu}^{2+}$ (34) | ESMBind-DL | **0.636** | **0.738** | **0.683** | **0.976** | **0.722** |
|  | M-Ionic | 0.530 | 0.630 | 0.576 | 0.955 | 0.611 |

**Table 2.** Summary of the protein-metal binding dataset, showing the number of sequences, the number of binding and non-binding residues for each metal type, and the percentage of binding residues relative to the total number of residues.

| Metal Ion | # Sequences | # Binding Residues | # Non-Binding Residues | % Binding Residues |
| --- | --- | --- | --- | --- |
| Zn<sup>2+</sup> | 2,101 | 9,614 | 629,724 | 1.53 |
| Ca<sup>2+</sup> | 1,809 | 9,543 | 633,812 | 1.51 |
| Mg<sup>2+</sup> | 2,033 | 6,582 | 720,702 | 0.91 |
| Mn<sup>2+</sup> | 602 | 2,347 | 212,024 | 1.11 |
| Fe<sup>3+</sup> | 291 | 1,231 | 94,398 | 1.30 |
| Co<sup>2+</sup> | 255 | 957 | 71,610 | 1.34 |
| Cu<sup>2+</sup> | 171 | 630 | 48,630 | 1.30 |

### Atomic Level Modeling

Transitioning from residue-level predictions to accurate 3D metal placement presents a significant challenge. The deep-learning model predicts high probabilities for most transition metals, complicating the determination of the specific metal type. To simplify this issue, we consider only one metal type per protein. Additionally, although ESMBind-DL predicts binding residues accurately, it doesn’t predict the number of metals present in the structure. We thus devised an effective metal placement strategy to tackle these challenges, optimizing metal positions based on geometric and energetic considerations (see Materials and Methods for details). To evaluate the performance of our method in predicting metal binding sites at the atomic level, we first define what constitutes a correct prediction. In our evaluation, a true positive is defined as a predicted metal within 5 Å of an experimental metal position in the protein structure. It’s important to note that this definition differs from the residue-level predictions discussed earlier; here, we are concerned about the accuracy of the 3D positions of the metals. We use several metrics to assess the performance of our method: precision, recall, F-1 score, and median deviation. Precision measures the proportion of correctly predicted metal sites among all predictions, while recall indicates the proportion of actual metal sites that were correctly identified. The F-1 score is the harmonic mean of precision and recall, providing a balanced measure of the two. The median deviation quantifies the spatial accuracy of the correctly predicted metal positions.

Figure 2(a) illustrates the precision-recall trade-off for each metal type. ESMBind demonstrates superior performance across various metal types, particularly for transition metals. For Zn^2+^, we achieve a high precision of 0.79 and a recall of 0.60, resulting in an F-1 score of 0.68. This indicates that our method is particularly effective at identifying true zinc-binding proteins and sites while minimizing false positives. Similarly, for Mn^2+^ and Fe^3+^, we obtain high precision (0.78 and 0.69 respectively) with good recall (0.58 and 0.64 respectively), resulting in F-1 scores of 0.67 and 0.66. The performance for lighter metals (element number < 20), Ca^2+^ and Mg^2+^, is somewhat lower, with a precision of 0.56 and 0.53, and recall of 0.31 and 0.32 respectively, yielding F-1 scores of 0.40 for both. This lower performance might be attributed to the greater variability in binding sites for Ca^2+^ and Mg^2+^ for their weaker and transient interactions compared to transition metals. The comparison of F-1 scores with Metal3D and AlphaFill is shown in Figure S1. To assess the overall performance across all metal types, we calculated the micro-averaged F-1 score, which gives equal weight to each prediction. ESMBind achieves a micro F-1 score of 0.54, outperforming Metal3D (0.51) and AlphaFill (0.46). This analysis demonstrates the robust overall performance of our method across different metal types.

**Figure 2.**
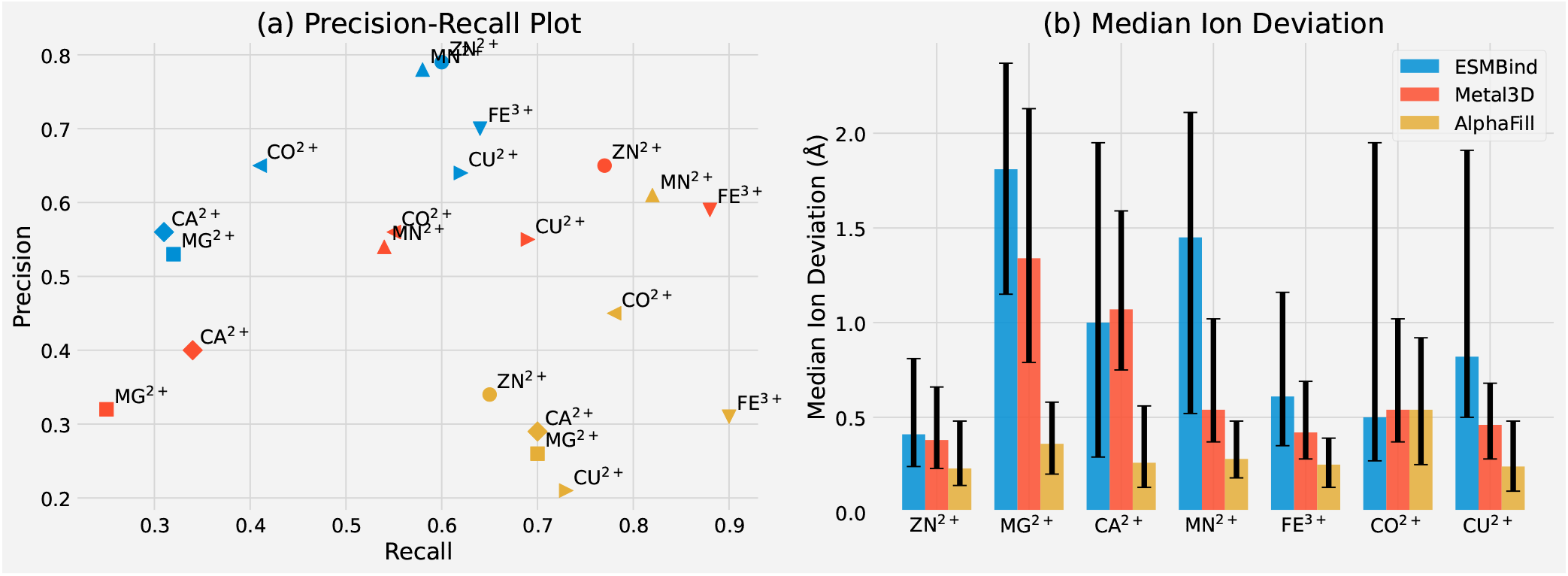
Performance comparison of ESMBind, Metal3D, and AlphaFill across various metal ions. (a) Precision-Recall plot. Different shapes represent different metal types. Different colors distinguish different methods. (b) Median deviation of predicted ion positions from their true locations. Error bars represent the 25th and 75th percentiles.

Figure 2(b) presents the median deviation of correctly predicted ion positions from their true locations. ESMBind generally shows higher median deviations compared to Metal3D and AlphaFill, which is expected given that our approach predicts the initial metal positions at the residue level instead of 3D structures. This intermediate step introduces a certain loss in spatial precision. However, it’s crucial to emphasize that these median deviations are calculated only for true positive predictions. When comparing ESMBind to Metal3D and AlphaFill, we observe distinct performance characteristics. ESMBind generally achieves higher precision across metal types, while AlphaFill tends to have higher recall but much lower precision. Metal3D often falls between the two in terms of precision and recall. The balance between precision and recall, as reflected in the F-1 score, is a key metric for evaluation. In this regard, ESMBind demonstrates competitive performance, achieving the highest micro-averaged F-1 score. This balanced performance highlights ESMBind’s robustness in handling diverse metal-binding scenarios, even though it may have slightly lower spatial accuracy for true positive predictions compared to structure-based methods.

### Ablation study

We conducted ablation studies to evaluate two key aspects of ESMBind: the deep learning model architecture (ESMBind-DL) and the ion placement strategy.

### Model architecture components

We assessed the contribution of different components in ESMBind-DL through systematic ablation experiments (Figure 3). The full model was compared with three variants: using only ESM-2 embeddings, using only ESM-IF embeddings, and a non-ensemble approach. The full model consistently outperformed all variants across seven metals, demonstrating the effectiveness of combining both sequence and structural information with an ensemble approach. The ESM-2-only model showed strong performance, particularly for transition metals, indicating that sequence information alone captures significant binding signals. In contrast, the ESM-IF-only model showed markedly decreased performance, suggesting that structural information alone is insufficient for accurate prediction. The non-ensemble model performed close to the full model, indicating that while ensembling improves robustness, the model’s primary strength comes from the integration of sequence and structural information.

**Figure 3.**
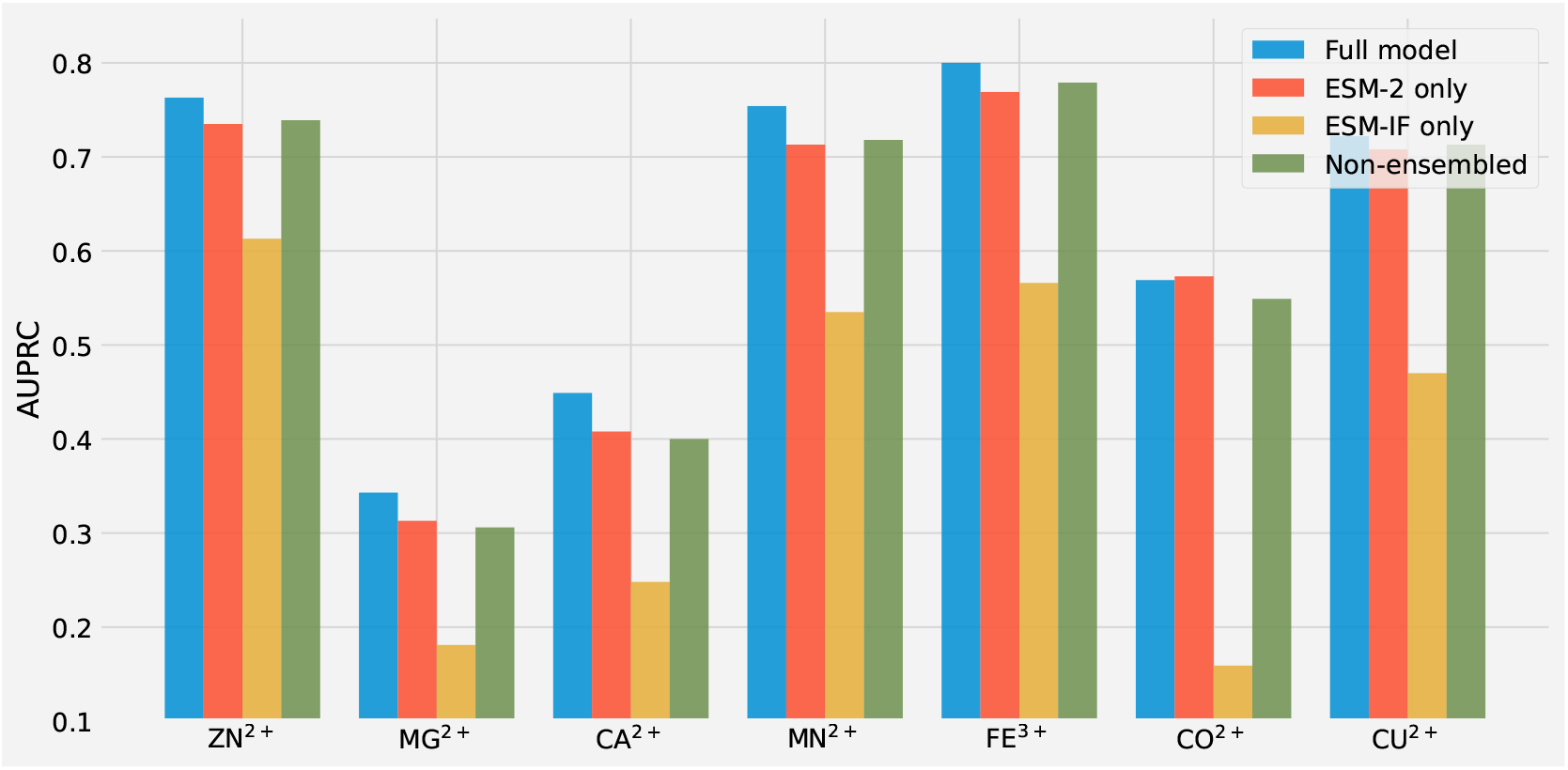
Ablation study of ESMBind-DL. Comparison of AUPRC scores between the full model and three variants: ESM-2 embeddings only, ESM-IF embeddings only, and non-ensemble approach. Results are shown for all seven metal types studied.

### Ion placement refinement

As illustrated in Figure 1(a), our workflow consists of three main stages: residue-level prediction, initial metal placement, and energy minimization for final refinement. To assess the contribution of the energy minimization stage, we compared the performance of the complete workflow against a version that omits this step (Table S1). Overall, energy minimization provides a modest performance improvement, increasing the micro-averaged F1 score from 0.535 to 0.538. The improvement was particularly notable for metals with more variable coordination geometries such as Mg^2+^ and Ca^2+^. For instance, in Mg^2+^ prediction, energy minimization increased true positives from 161 to 167 while maintaining similar false positive rates. This suggests that energy minimization is most beneficial for refining less accurate initial predictions and may be particularly valuable when applied to predicted protein structures, where the minimization step can help ensure predictions are energetically favorable.

### Predict novel metal-binding proteins and structures

To further evaluate the utility of the workflow in predicting metal binding proteins and structures, we applied it to 142 putative effector proteins secreted by *Colletotrichum sublineola* (*C. sublineola*), a fungal pathogen leading to Sorghum anthracnose^16, 17^. These putative effector proteins could modulate the plant’s immune system for pathogen’s virulence and infection. However, none of their functions have been predicted and characterized. Using the sequences and Alphafold models of the 142 effectors as input to our workflow, we predict that, out of 142, 22 bind to Zn^2+^, 18 bind to Mn^2+^, and 21 bind to Fe^3+^ (Table S2). We used the pLDDT scores from the AlphaFold-predicted models to rank the predicted metal-binding proteins and presented the top four in Figure 4. The highest pLDDT score is a zinc-finger protein (Uniprot# A0A066X8W9). A Zn^2+^ is coordinated by four cysteine residues (Figure 4a). A functional implication could be that this effector protein might modulate its host’s RNA/DNA stability or activity in favor of infection and virulence. The prediction of zinc-finger protein is supported by a yeast homolog structure (PDB id 2J6A) of a Trm112 protein, which is a methyltransferase activator^18^. The superimposition of the two structures has an RMSD of only 0.87 Å with an almost identical Zn^2+^ location (Figure S3). Similarly, we predicted A0A066XCS5 is an iron-binding, Hce2 domain-containing effector protein^19^, A0A066XI99 is a manganese-binding, mannosylphosphate transferase ^20^, and A0A066XXV0 is a lysine-specific metalloendopeptidase domain-containing protein ^21^ (Figure 4b,c,d).

**Figure 4.**
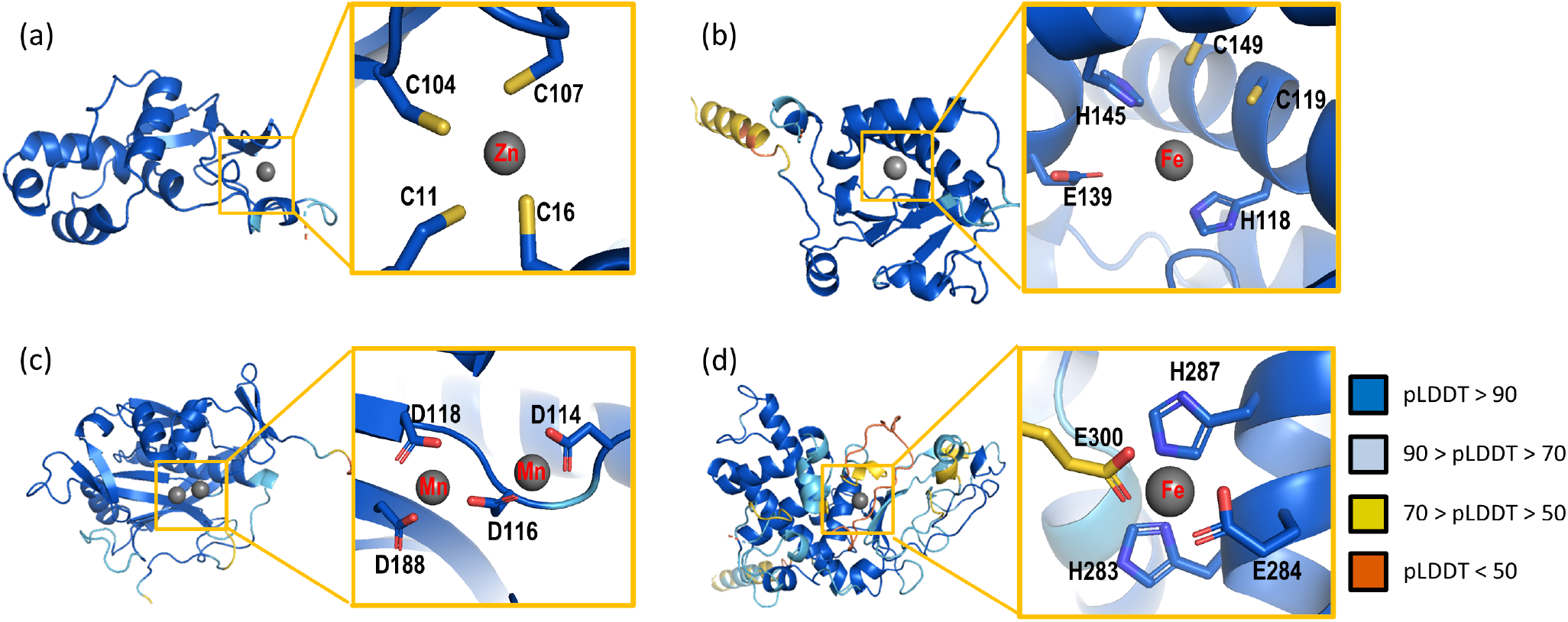
Predicted metal binding proteins and structures. The top four results from the application of ESMBind workflow to purtative C. sublineola effectors colored by pLDDT scores with metal ions shown in grey. (a) A0A066X8W9 (pLDDT 96.2) is predicted to be a Trm112p-like protein. (b) A0A066XCS5 (pLDDT 86.83) is predicted to be a Hce2 domain-containing effector protein (c) A0A066XI99 (pLDDT 85.5) is predicted to be a mannosylphosphate transferase. (d) A0A066XXV0 (pLDDT 83.58) is predicted to be a lysine-specific metallo-endopeptidase domain-containing protein.

## Discussion

The performance of ESMBind demonstrates the effectiveness of integrating foundation models with physics-based approaches for metal binding prediction. Our analysis highlights several key aspects of the method’s capabilities and limitations:

### Architecture and model design

As shown in Figure 1(b), ESMBind-DL combines sequence and structure information through ESM-2 and ESM-IF, respectively. Our ablation studies (Figure 3) demonstrate that this dual-input strategy consistently outperforms single-input approaches. The success of our simple MLP architecture over the Transformer-based architectures of LMetalsite and M-Ionic (Table 1) suggests that the integration of both sequence and structure information provides sufficiently rich features that can be effectively leveraged without complex downstream processing. A practical limitation to consider is the computational requirement of two pre-trained models, particularly ESM-IF’s need for protein structure input. However, as demonstrated in our case study with *C. sublineola* effectors (Figure 4), AlphaFold-predicted models can serve as effective inputs since ESM-IF has been trained on an extensive set of AlphaFold structures.

### Metal placement strategy

Our metal placement strategy effectively bridges residue-level predictions and atomic-level 3D metal-binding sites. The benchmarking results (Figure 2) show that ESMBind achieves higher precision and recall compared to Metal3D and AlphaFill, though with slightly lower spatial accuracy. This trade-off reflects our two-stage approach versus Metal3D’s end-to-end 3D prediction or AlphaFill’s direct transplant method. For many applications, particularly when predicting metal-binding in proteins with low sequence homology to experimental structures, accurate metal binding site identification outweighs small differences in spatial precision. The effectiveness of this approach is evident in our successful prediction of novel metal-binding proteins in *C. sublineola* (Figure 4), where we identified previously unknown metal-binding functions in putative effector proteins.

### Limitations

While ESMBind’s training data (Table 2) comes from BioLip’s curated dataset of biologically relevant binding sites, protein structure databases may contain misidentified metals, as documented by Refs.^2, 22, 23^. Beyond data quality considerations, a fundamental limitation of ESMBind is that it currently predicts binding sites for each metal ion independently rather than determining metal selectivity. While the model outputs binding probabilities for multiple metals per residue, these should not be interpreted as predictions of preferential binding. This design choice reflects the complex nature of metal selectivity in proteins, which depends on multiple factors beyond local binding site architecture, including protein matrix effects, cellular metal homeostasis, and metal ion concentrations^24^. True metal selectivity prediction would require integration of broader protein structural context, dynamic binding aspects, and cellular metal trafficking considerations. Nevertheless, our current approach effectively identifies geometrically and chemically compatible metal binding sites to guide experimental characterization.

### Future directions

ESMBind balances identification accuracy with reasonable spatial precision, demonstrating the potential of hybrid approaches that integrate deep learning with physics-based modeling. Future work could focus on improving spatial accuracy through more sophisticated energy models or exploring end-to-end learning approaches that maintain the benefits of pre-trained foundation models. Beyond metal ions, our framework could be extended to predict binding sites for other types of ligands. The combination of sequence-based and structure-based foundation models, coupled with physics-based refinement, provides a general recipe for predicting protein-ligand interactions. Such extensions could be particularly valuable for drug discovery and protein engineering applications where accurate prediction of diverse ligand binding sites is crucial.

## Materials and methods

### Dataset collection and process

We retrieved protein-metal binding data from the BioLip database^14, 25^. BioLip parses protein structures from the Protein Data Bank (PDB) to identify biologically relevant ligands and their binding residues. For any biologically relevant ligand, it defines a residue as binding if the closest atomic distance between the residue and the ligand is within 0.5 Å plus the sum of Van der Waal’s radii of the two atoms.

We downloaded data from BioLip on 05-30-2024. We filtered the data to only include sequences with a maximum length of 1500 residues and structures with an experimental resolution of 3.0 Å or better. To obtain a non-redundant set of sequences for each metal type, we used CD-HIT^26^ with a 30% sequence identity cutoff. Finally, we randomly split the resulting sequences into training (80%) and test (20%) sets. The test dataset was reserved for evaluating the trained models. We compiled a summary of the dataset, detailing the number of sequences, binding residues, and non-binding residues for each metal ion, as shown in Table 2.

### Model training

The training process for ESMBind involves several key components and techniques to optimize performance and handle the challenges associated with the highly imbalanced dataset. We trained the model on multiple metal types simultaneously, with separate train and validation datasets prepared for each metal type, using the binary cross-entropy loss as the loss function. To address the issue of imbalanced data, where the number of negative samples (non-binding residues) greatly exceeds the number of positive samples (binding residues), we employ an imbalanced dataset sampler that adjusts the sampling probability of each class based on the desired positive ratio. Specifically, we boost the probability of the positive class to 3-fold of its original ratio, effectively oversampling the minority class. During training, we evaluate the model’s performance using the Area Under the Precision-Recall Curve (AUPRC) metric, which is particularly suitable for evaluating imbalanced datasets. We also determined the optimal classification threshold for each metal type by finding the threshold that maximizes the F1 score on the validation set. We stored the best threshold for each metal type to use with binary predictions.

We employ regularization techniques at both the model and training levels to prevent overfitting and improve the model’s robustness. At the model level, we apply dropout and layer normalization and inject Gaussian noise into the embeddings during training. At the training level, we use weight decay as a regularization term in the optimizer and implement early stopping by monitoring the validation AUPRC, stopping training if there is no improvement after 7 epochs, and saving the model with the best validation AUPRC as the final model. We trained the model using the AdamW optimizer with a weight decay of 0.035 and employed a learning rate scheduler to adjust the learning rates during training by reducing the learning rate by a factor of 0.7 every 15 epochs. This helped the model converge to a better solution by taking smaller steps as the training progressed. Additionally, we applied Stochastic Weight Averaging^27^ (SWA) to further improve generalization and stability, where we updated a separate SWA model with the average of the model parameters after a specified number of epochs, which acts as another regularization technique.

To further boost the performance, we employed an ensemble approach. We split the entire dataset into an 80:20 ratio, reserving the 20% split as the final test dataset. We divide the remaining 80% of the data into 5 folds for cross-validation and train each model in the ensemble on 4 folds while validating on the remaining fold. The final ensemble model comprises 5 models, each trained on a different combination of training and validation folds. During inference, we average the predictions from these 5 models to obtain the final output probabilities. This ensemble approach reduces overfitting and improves the robustness and generalization ability of the model.

### Metal placement and energy minimization

Building upon the residue-level binding predictions from ESMBind, we developed a protocol to place metals in 3D protein structures and optimize their positions. This process involves several key steps:

### Initial Metal Placement

We use the binding probabilities output by ESMBind to identify potential binding residues. To determine binding residues, we employed a threshold set at half of the optimal threshold obtained during model validation. This optimal threshold is defined as the threshold that maximizes the F1 score on the validation set during the ESMBind training process. By using half of this optimal threshold, we address the model’s tendency to produce more false negatives than false positives due to the highly imbalanced dataset distribution. For each predicted binding residue, we identified all potential binding atoms based on the metal type and then calculated the center of mass of these binding atoms to determine the initial metal position. This approach ensures that the initial metal placement considers all relevant binding atoms for each residue. The list of potential binding atoms for different residues is based on the study by Zheng et al.^28^ and Bazayeva et al.^2^.

### Binding Site Identification

We employed agglomerative hierarchical clustering to group nearby binding metals after their initial placement. The process uses a dynamic distance threshold, starting with a predefined value (7Å) and gradually decreasing it until at least one cluster is identified. For each resulting cluster, we merge the binding atoms from all constituent placements and calculate a new metal position as the center of mass of these merged atoms. This approach allows for the consolidation of multiple potential binding sites into distinct metal ion locations.

### Refined Metal Positioning

We implemented a local grid search strategy to optimize the metal positions. This approach aims to place the predicted metals as close to their binding protein atoms as possible while avoiding steric clashes. The search considers ideal binding distances for different atom types and evaluates potential positions based on a scoring function that balances favorable interactions with binding atoms and unfavorable close contacts with non-binding atoms. This method allows for fine-tuning of metal positions within the local environment of the binding site.

### System Construction

We constructed the molecular system using the AMBER force field^29^, incorporating a single protein chain, one metal type, and water as the solvent. The 12-6-4 Lennard-Jones non-bonded model^30^ characterizes metal interactions, providing an accurate representation of the metal-protein and metal-solvent interactions.

### Energy Minimization

We used OpenMM^31^ to perform a series of constrained energy minimizations: a) Apply restraints to all non-solvent atoms, allowing water molecules to adapt to the protein-metal complex. b) Restrain protein backbone atoms and metals, permitting side chain adjustment. c) Constrain only metals, enabling protein relaxation around optimized metal locations. The force constant is halved with each step, facilitating gradual structural adaptation. This multi-step minimization process allows for the optimization of the metal positions while maintaining the overall protein structure.

This protocol leverages the strengths of deep learning for initial binding site prediction, employs a local search algorithm to refine metal locations, and uses energy minimization to further optimize the protein-metal complex. By addressing the challenges of transitioning from residue-level predictions to precise 3D ion placement, our method provides a robust and physically realistic approach to identifying and characterizing metal-binding proteins and structures.

### Prediction of metal-binding pathogen effectors

The proteome of sorghum anthracnose fungal pathogen *Colletotrichum sublineola* (*C. sublineola*) was retrieved from Uniprot (www.uniprot.org). The proteome contains 12687 unique protein sequences. We used the program SignalP 6^32^ to predict signal peptides, WolF-PSORT^33^ to identify their subcellular localizations, and we obtained 1370 targets as secreted proteins. Using the effector prediction program EffectorP^34^, we predicted and selected 142 putative effectors that are secreted by *C. sublineola* and function in the host’s cytoplasm. We used their AlphaFold structure models and sequences as input for the prediction of metal-binding proteins and modeling of the protein-metal complexes.

## Acknowledgments

We thank Kriti Chopra for the helpful discussion. This work was supported in part by Brookhaven National Laboratory LDRD 23-06 and the U.S. Department of Energy (DOE), Office of Biological and Environmental Research (KP1601011).

## Data and code availability

This study utilizes publicly available data from three main sources: protein structures from the Protein Data Bank (https://www.rcsb.org), AlphaFold predicted structures (https://alphafold.ebi.ac.uk/), and binding data from BioLiP (https://zhanggroup.org/BioLiP/index.cgi). ESMBind is developed as an open-source Python package and is freely available on GitHub (https://github.com/Structurebiology-BNL/ESMBind). This repository contains all the necessary code for both training and testing the model.

