## Supplemental Materials for "Predicting metal-binding proteins and structures through integration of evolutionary-scale and physics-based modeling"

Table S1: Impact of energy minimization on metal placement performance. The table shows true positives (TP), false positives (FP), and false negatives (FN) for each metal type, comparing results with and without the energy minimization step.

| Metal | With Energy Minimization | | | Without Energy Minimization | | |
|---|---|---|---|---|---|---|
| | TP | FP | FN | TP | FP | FN |
| $Zn^{2+}$ | 379 | 100 | 250 | 377 | 97 | 249 |
| $Ca^{2+}$ | 168 | 132 | 369 | 166 | 131 | 371 |
| $Mg^{2+}$ | 167 | 150 | 352 | 161 | 146 | 347 |
| $Mn^{2+}$ | 109 | 31 | 76 | 106 | 30 | 79 |
| $Fe^{3+}$ | 46 | 20 | 26 | 44 | 16 | 28 |
| $Co^{2+}$ | 26 | 14 | 38 | 25 | 14 | 39 |
| $Cu^{2+}$ | 34 | 19 | 20 | 34 | 19 | 20 |
| Micro-averaged F1: 0.538 (with) vs 0.535 (without) | | | | | | |

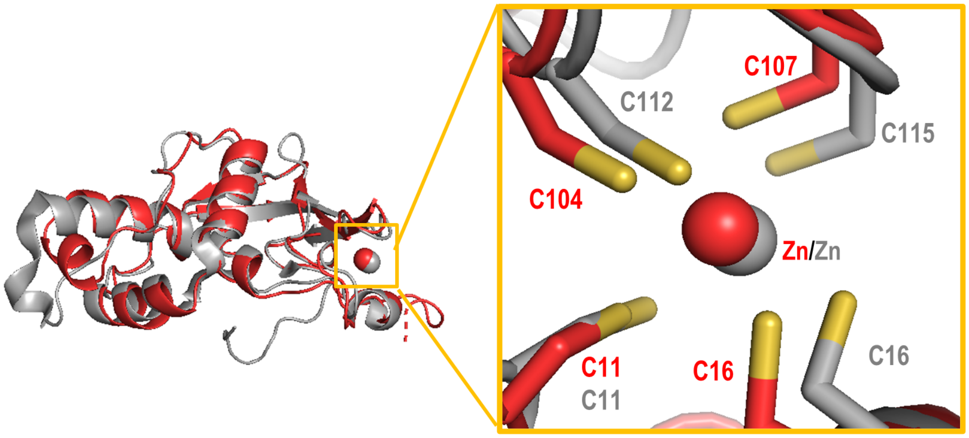


Figure S1: Comparison of a predicted fungal metal binding protein structures with its experimental homolog structures in PDB. C. sublineola A0A066X8W9 (red), a Trm112p-like protein bound with Zn aligned with S. cerevisiae Trm112 protein (PDB code 2J6A, grey) which is a methyltransferase activator.

Table S2: Metal binding predictions for *C. sublineola* effector proteins (Part 1/3)

| **UniProt ID** | $\mathbf{Zn^{2+}}$ | $\mathbf{Mn^{2+}}$ | $\mathbf{Fe^{3+}}$ |
|---|---|---|---|
| A0A066WSI5 | • | | |
| A0A066WTH2 | • | • | • |
| A0A066WTT5 | | | |
| A0A066WUF0 | | | |
| A0A066WUN1 | | | |
| A0A066WV46 | | | |
| A0A066WV97 | | | |
| A0A066WVP9 | | | |
| A0A066WX93 | • | • | • |
| A0A066WXB0 | | | |
| A0A066WXE5 | | | |
| A0A066WXG4 | | | |
| A0A066WXH2 | | | |
| A0A066WXV8 | • | • | • |
| A0A066WYT5 | | | |
| A0A066WZ31 | | | |
| A0A066WZF9 | | | |
| A0A066WZH1 | | | |
| A0A066WZI1 | | | |
| A0A066WZR3 | | | |
| A0A066X0P5 | | • | • |
| A0A066X110 | | | |
| A0A066X1N3 | | | |
| A0A066X254 | | | • |
| A0A066X294 | | | |
| A0A066X2B8 | | | |
| A0A066X2G9 | | | |
| A0A066X2J5 | • | • | • |
| A0A066X2M6 | | | |
| A0A066X320 | | | |
| A0A066X3P1 | | | |
| A0A066X480 | | | |
| A0A066X4J0 | | | |
| A0A066X4N4 | | | |
| A0A066X4U8 | • | • | • |
| A0A066X4Y3 | • | • | • |
| A0A066X634 | | | |
| A0A066X6T9 | | | |
| A0A066X6Y7 | | | |
| A0A066X779 | | | |
| A0A066X7S1 | | | |
| A0A066X7Y3 | | | |
| A0A066X825 | | | |
| A0A066X835 | • | • | • |
| A0A066X8J7 | | | |
| A0A066X8N1 | | | |
| A0A066X8S2 | | | |
| A0A066X8W9 | • | | • |
| A0A066X951 | | | |
| A0A066X9B2 | | | |

Table S3: Metal binding predictions for *C. sublineola* effector proteins (Part 2/3)

| **UniProt ID** | $\mathbf{Zn^{2+}}$ | $\mathbf{Mn^{2+}}$ | $\mathbf{Fe^{3+}}$ |
|---|---|---|---|
| A0A066X9H2 | | | |
| A0A066X9K0 | | | |
| A0A066X9Q0 | | | |
| A0A066X9W4 | | | |
| A0A066XA99 | | | |
| A0A066XAT9 | | | |
| A0A066XB77 | | | |
| A0A066XBD8 | | | |
| A0A066XBF9 | | | |
| A0A066XBH5 | | | |
| A0A066XBI3 | • | | • |
| A0A066XBK1 | | | |
| A0A066XBL9 | | | |
| A0A066XBR8 | | | |
| A0A066XC28 | | | |
| A0A066XCA8 | | | |
| A0A066XCK7 | | | |
| A0A066XCS5 | • | • | • |
| A0A066XD89 | | | |
| A0A066XE86 | | | |
| A0A066XE94 | | | |
| A0A066XEG5 | | | |
| A0A066XET4 | | | |
| A0A066XEU7 | | | |
| A0A066XFQ9 | | | |
| A0A066XFR2 | • | • | • |
| A0A066XFT1 | | | |
| A0A066XGG6 | | | |
| A0A066XH10 | | | |
| A0A066XH13 | | | |
| A0A066XH53 | | | |
| A0A066XH92 | | | |
| A0A066XHE8 | • | • | • |
| A0A066XHQ0 | | | |
| A0A066XI99 | • | • | • |
| A0A066XIB3 | | | |
| A0A066XID5 | | | |
| A0A066XIF4 | | | |
| A0A066XIU8 | • | | |
| A0A066XIY2 | | | |
| A0A066XJQ7 | | | |
| A0A066XK85 | | | |
| A0A066XL46 | | | |
| A0A066XLH0 | | | |
| A0A066XLM2 | | | |
| A0A066XLM6 | | | |
| A0A066XLS9 | | | |
| A0A066XLT5 | | | |
| A0A066XLV3 | • | | |
| A0A066XM58 | | | |

Table S4: Metal binding predictions for *C. sublineola* effector proteins (Part 3/3)

| **UniProt ID** | **$Zn^{2+}$** | **$Mn^{2+}$** | **$Fe^{3+}$** |
|---|---|---|---|
| A0A066XM65 | | • | • |
| A0A066XM92 | • | | |
| A0A066XMH0 | | | |
| A0A066XN61 | | | |
| A0A066XP44 | | | |
| A0A066XPS5 | | | |
| A0A066XPY2 | | | |
| A0A066XPZ2 | | | |
| A0A066XQE2 | | | |
| A0A066XQF2 | | | |
| A0A066XQK3 | | | |
| A0A066XQS2 | | | |
| A0A066XQZ8 | | • | • |
| A0A066XRK6 | • | • | • |
| A0A066XRT4 | | | |
| A0A066XRT8 | | | |
| A0A066XRW4 | | | |
| A0A066XRZ1 | | | |
| A0A066XS45 | | | |
| A0A066XSA9 | | | |
| A0A066XSI2 | | | |
| A0A066XSM0 | | | |
| A0A066XSV3 | • | • | • |
| A0A066XSV7 | | | |
| A0A066XSZ6 | | | |
| A0A066XT41 | | | |
| A0A066XTC3 | | | |
| A0A066XTJ4 | • | | |
| A0A066XVL0 | | • | • |
| A0A066XVM7 | | | |
| A0A066XW89 | | | |
| A0A066XWB7 | | | |
| A0A066XWE7 | | | |
| A0A066XWP2 | | | |
| A0A066XXI5 | • | | |
| A0A066XXV0 | • | • | • |
| A0A066XZD4 | | | |
| A0A066XZR5 | | | |
| A0A066Y1B0 | | | |
| A0A066Y1J3 | | | |
| A0A066Y2D0 | | | |
| A0A066Y2E5 | | | |
